# Oligodendrocyte precursor cells engulf synaptic inputs in an experience- and microglia-dependent manner

**DOI:** 10.1101/2022.02.10.479887

**Authors:** Yohan S.S. Auguste, Austin Ferro, Jessica Dixon, Uma Vrudhula, Jessica Kahng, Anne-Sarah Nichitiu, Lucas Cheadle

## Abstract

Oligodendrocyte precursor cells (OPCs) are a highly proliferative class of non-neuronal progenitors that largely give rise to myelinating oligodendrocytes. Although OPCs persist across the lifespan, their functions beyond oligodendrogenesis remain to be fully characterized. Here, we show that OPCs contribute to neural circuit remodeling by internalizing presynaptic thalamocortical inputs in both the developing and adult mouse visual cortex. Inputs internalized by OPCs localize to lysosomal compartments, consistent with OPC engulfment of synapses occuring through phagocytosis. We further show that engulfment by OPCs is heightened during experience-dependent plasticity, and that this experience-dependent increase in engulfment requires microglia. These data identify a new function for OPCs beyond the generation of oligodendrocytes and reveal that distinct non-neuronal populations collaborate to modulate synaptic connectivity.

## Manuscript

The refinement of synapses in response to sensory experience sculpts brain connectivity during late stages of postnatal development and facilitates neural circuit plasticity in the adult^1^. However, the cellular and molecular mechanisms underlying experience-dependent refinement remain poorly understood. Recent work demonstrates that microglia and astrocytes, prominent populations of non-neuronal brain cells collectively called glia, promote synaptic refinement in the mammalian visual system by eliminating excess synapses prior to the onset of sensory experience at eye-opening, which occurs around postnatal days (P)12 – P14^2,3^. Astrocytes and microglia predominantly eliminate synapses during this time by phagocytosing (i.e. eating) synaptic components and digesting them within acidic lysosomal compartments. While these paradigm-shifting discoveries unveiled synapse engulfment by glia as a core biological mechanism through which non-neuronal cells shape circuit connectivity and function early in life, the possibility that glia serve as intermediaries between visual experience and synaptic refinement during late stages of development and in the adult remained to be extensively tested.

<u>O</u>ligodendrocyte <u>p</u>recursor <u>c</u>ells (OPCs) are a specialized population of glial progenitors in the brain that give rise to myelinating oligodendrocytes, cells that promote communication between neurons by wrapping their axons in myelin sheaths^4^. Initially born in the subventricular zones of the embryonic neural tube, OPCs migrate throughout the brain and spinal cord where they continue to proliferate and differentiate into oligodendrocytes, and less frequently other cell types, well into postnatal development^5,6^. Although the rate of oligodendrocyte production by OPCs decreases significantly as the brain matures, OPCs remain abundant and maintain their multipotent capacity in the adult brain. OPCs also dynamically respond to a variety of cues, including neuronal activity and molecular signals from microglia, by differentiating into oligodendrocytes and thereby increasing the production of myelin^7,8^. OPCs also receive direct synaptic input from neurons, though the functions of OPC:neuron synapses remain to be defined^9,10^. In addition to their ability to form synaptic connections, the persistence of OPCs across the lifespan even after mature myelination patterns have been established suggests that OPCs may play important roles in the brain beyond their contributions as an adult progenitor pool. However, such alternative functions have yet to be extensively characterized.

To uncover experience-dependent mechanisms of synaptic refinement in the postnatal brain, we analyzed interactions between non-neuronal cells including OPCs and presynaptic thalamocortical (TC) inputs from the <u>d</u>orsal <u>l</u>ateral <u>g</u>eniculate <u>n</u>ucleus (dLGN) of the thalamus as they synapse onto their postsynaptic targets in layer 4 of the primary visual cortex (V1) of the mouse. We chose the visual TC circuit as the basis for this study because it undergoes a well-defined period of heightened experience-dependent synaptic development during the third week of life, and because synapse elimination in the adult contributes to vision-dependent plasticity^11,12^ (Fig. 1A). We first assessed interactions between glia and synapses across postnatal development and in the mature brain by immunostaining for proteins enriched in distinct non-neuronal cell types along with the TC input marker Vglut2 at postnatal days (P)10 (prior to eye-opening), P20 (the beginning of the critical period of sensory-dependent remodeling), P27 (the peak of sensory-dependent remodeling), and P90, when the brain is fully mature. By quantifying these interactions using a well-established engulfment assay^13^, we identified TC inputs within microglia at all time points analyzed, confirming that microglia engulf synapses in V1 as the brain matures.

**Figure 1.**
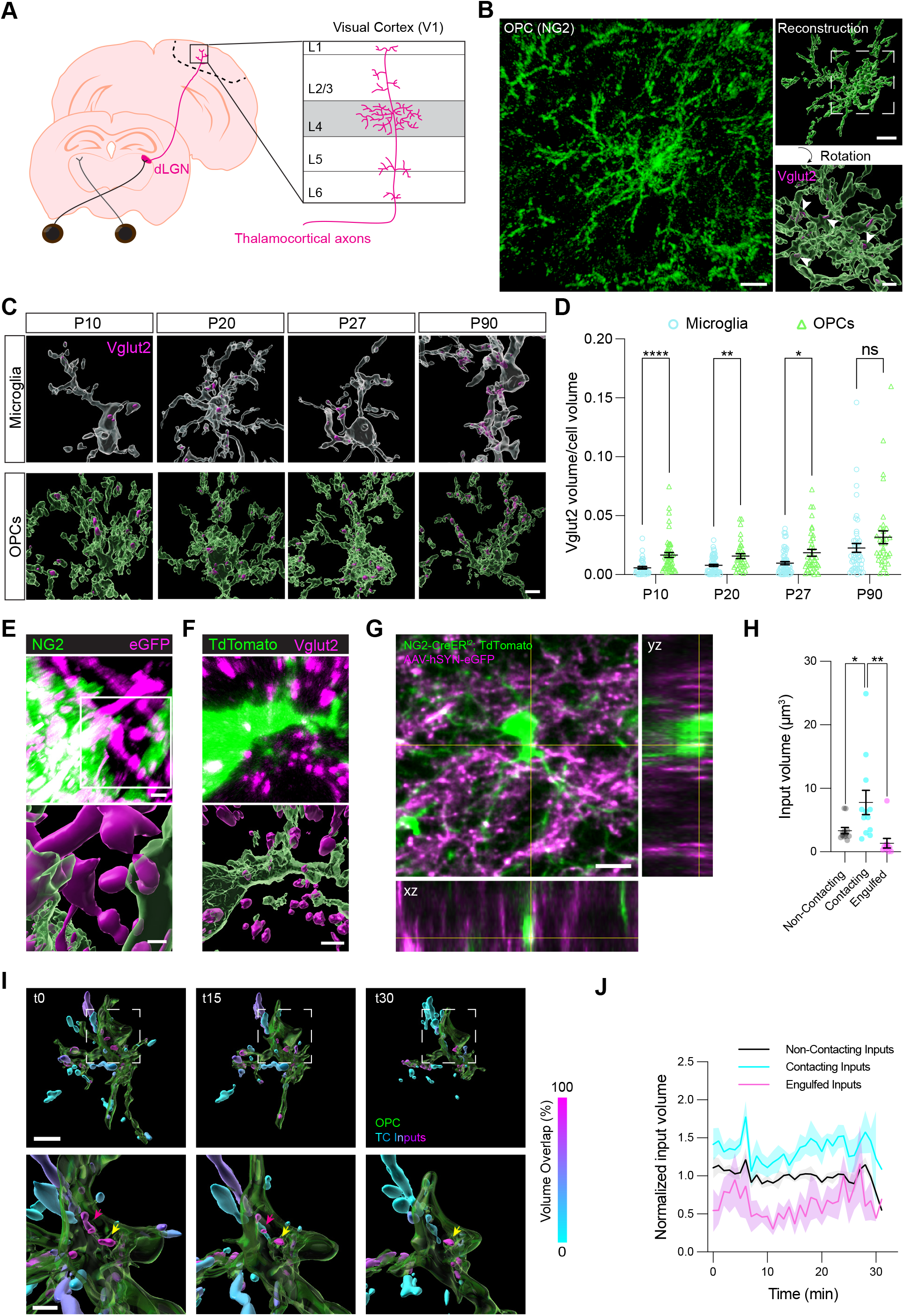
OPCs engulf thalamocortical synaptic inputs in primary visual cortex. (A) Schematic of thalamocortical (TC) inputs from the dorsal lateral geniculate nucleus (dLGN) synapsing onto layer four of primary visual cortex (V1). (B) Confocal image of an oligodendrocyte precursor cell (OPC) immunostained for NG2 (green) alongside volumetric reconstructions of the OPC with and without engulfed synaptic inputs immunostained for Vglut2 (magenta; white arrows). Scale bar, 10 μm. Inset scale bar, top: 5 μm; bottom: 2 μm. (C) Volumetric reconstructions of microglia (Iba1, white) and OPCs (NG2, green) containing TC inputs (Vglut2, magenta) at multiple postnatal ages. Scale bar, 5 μm. (D) Quantification of the volume of synaptic material contained within the microglial or OPC volume. Two-way ANOVA with Geisser-Greenhouse correction (Cell type: p < 0.0001; age: p < 0.0001; interaction between cell type and age: p > 0.05) and Šídák multiple comparisons. n (microglia/OPC): P10 = 38/53, P20 = 63/35, P27 = 60/39, P90 = 49/36, from 3 mice per group. (E) Confocal image and reconstruction of an NG2-immunostained OPC (green) containing AAV-hSYN-eGFP+ TC inputs from the dLGN pseudocolored in magenta. Scale bar, 2 μm. Inset scale bar, 2 μm. (F) Confocal image and reconstruction of an OPC in a NG2-CreER^T2^; TdTomato+ reporter mouse (pseudocolored in green) containing Vglut2-immunostained synaptic inputs (magenta). Scale bar, 2.5 μm. (G) Two-photon image of a TdTomato+ OPC (green) interacting with AAV-hSYN-eGFP+ TC axons and inputs (magenta) in V1 of an awake mouse. Orthogonal projections shown below and to the right. Scale bar, 10 μm. (H) Quantification of the average volumes of synaptic inputs depending upon whether they contact OPCs, are engulfed by OPCs, or do not interact with OPCs. One-way ANOVA (p < 0.0001) with Tukey post-hoc test. n = 12 volumes from 3 mice. (I) Volumetric reconstructions of OPCs (green) interacting with synaptic inputs colored based upon percentage overlap with the OPC (color legend to right of images). Inputs that are completely internalized are shown in magenta while those that do not contact the OPC are in cyan. Representative images are taken from a thirty-minute time-lapse imaging session shown in supplemental movie 1. Scale bars, top: 10 μm; bottom: 5 μm. Yellow arrows, engulfed input that is present throughout imaging session. Magenta arrows, engulfed input that disappears during session. (J) Average volume of all inputs across the imaging period depending upon whether the inputs are in contact with or engulfed by OPCs. Mixed effects analysis of synapse category, p < 0.001, time p < 0.05. n = 6 videos taken from 3 mice. In (D) and (H), individual data points shown with bars representing mean +/− SEM. *p< 0.05, **p< 0.01, ****p<0.0001. In (J), solid lines represent mean and shaded areas represent SEM.

Unexpectedly, although microglia are considered to be the primary phagocytes of the brain, we observed that OPCs not only contained TC inputs within their cellular boundaries, but that they contained more Vglut2+ inputs compared to microglia at all developmental time points analyzed, and the same level of inputs in the adult (Fig. 1B-D). Initially focusing on the P90 time point, we complemented the antibody-based method for labeling presynaptic terminals *ex vivo* by labeling TC inputs *in vivo* through infection of neurons in the dLGN with AAV9-hSYN-eGFP (Extended Data Fig. 1A). This experiment confirmed the presence of GFP+ structures within OPCs in V1 (Fig. 1E). Reciprocally, we labeled OPCs *in vivo* by crossing the NG2-CreER^T2^ mouse line^14^ with the Lox-STOP-Lox-TdTomato reporter line (NG2-CreER^T2^; TdTomato mice; Extended Data Fig. 1B-F), and immunostained V1 sections from these mice for Vglut2. This analysis confirmed the presence of Vglut2+ TC inputs within the boundaries of TdTomato+ OPCs (Fig. 1F).

Next, to determine whether OPCs express the molecular machinery necessary to engulf synapses, we assessed the expression of a number of genes encoding known mediators of phagocytosis and lysosomal function both by re-analyzing a published single-cell RNA-sequencing dataset from adult mouse V1^15^ and through multiplexed fluorescence *in situ* hybridization (RNAscope). Relevant genes were taken from the “phagocytosis” gene ontology term from the PANTHER database^16^. These analyses confirmed that OPCs express molecular regulators of engulfment, including *Ptprj, Calcrl, Mertk,* and *Arsb* (Extended Data Figure 2). Altogether, these data provide evidence that OPCs engulf presynaptic terminals in the mammalian neocortex.

Because imaging interactions between OPCs and synapses in sections *ex vivo* requires tissue fixation and permeabilization which can sometimes obscure structural interactions, we next sought to visualize synapses within OPCs in the living brain. Toward this end we again infected the dLGNs of adult NG2-CreER^T2^; TdTomato mice with AAV9-hSYN-eGFP then imaged interactions between OPCs and TC inputs in layers 3 and 4 of V1 in awake mice by two-photon microscopy (Extended Data Fig. 3A). First, we captured single-timepoint volumes of V1 containing TdTomato+ OPCs interacting with GFP+ TC inputs, and quantified the number of synaptic inputs that interacted with each OPC in the field of view. Consistent with previous reports that OPCs receive synaptic input from neurons^9,10^, we found that every OPC was in direct contact with at least one eGFP+ TC input, and that each OPC interacted with 16.5 ± 2.85 inputs on average (Fig. 1G and Extended Data Fig. 3B-E). Furthermore, we observed that a large majority (88.5%) of OPCs contained eGFP+ material within their cellular boundaries.

To further explore interactions between OPCs and TC inputs, we next separated synaptic inputs into three categories based upon their proximity to the OPC surface: inputs that did not contact OPCs (non-contacting), inputs that contacted OPCs, and inputs that were internalized by OPCs (i.e. those that were at least 270 nm internal to the OPC surface). This analysis revealed that inputs in direct contact with the OPC surface were significantly larger than non-contacting inputs as well as internalized inputs (Fig. 1H), and that internalized inputs were smaller than both contacting and non-contacting terminals (Extended Data Fig. 3F). This observation is consistent with the engulfed inputs representing synaptic components that may be undergoing digestion.

To determine the short-term stability of inputs engulfed by OPCs, we next performed time-lapse volumetric imaging of interactions between TC inputs and OPCs over a 30-minute period of time in V1 of awake, unanesthetized mice (Supplemental Movies 1 and 2). These data revealed that inputs engulfed by OPCs remained small and relatively stable across this time frame, though in rare cases we observed an input that was initially within an OPC but disappeared during the imaging period, again suggesting the possibility that internalized inputs are being digested (Fig. 1I,J). These data suggest that OPCs engulf TC inputs *in vivo.*

Synaptic components engulfed through phagocytosis are shuttled into acidic lysosomal compartments for digestion, as evidenced by a strong colocalization of synaptic inputs inside of microglia with markers of lysosomal membranes^2^. To determine the fate of TC inputs engulfed by OPCs, we immunostained the brains of NG2-CreER^T2^; TdTomato mice for Vglut2 and the late-lysosomal membrane protein Lamp2. Using standard confocal microscopy, we observed a substantial population of engulfed inputs that resided within Lamp2+ lysosomes, consistent with OPCs eliminating inputs through engulfment, similar to microglia. We also observed a population of internalized inputs that did not colocalize with Lamp2 and may represent structures that are at different stages of endosomal processing, or that interact with OPCs through a non-endocytic mechanism (Fig. 2A). <u>S</u>uper-resolution <u>S</u>tructured <u>I</u>llumination <u>M</u>icroscopy (SIM) of V1 sections from NG2-CreER^T2^; TdTomato mice infected with AAV9-hSYN-eGFP and stained for Lamp2 confirmed the presence of engulfed synapses within OPC lysosomes (Fig. 2B).

**Figure 2.**
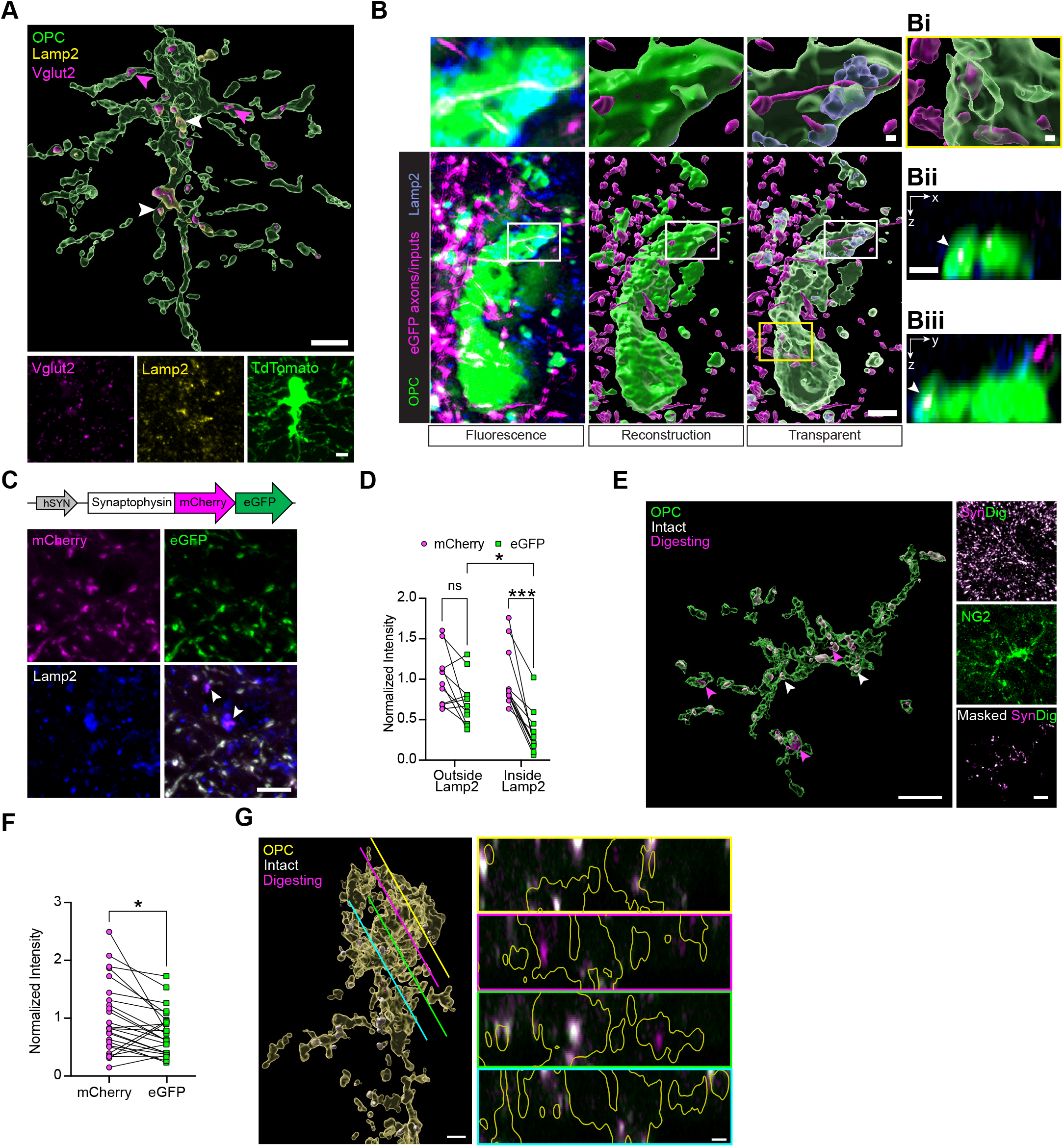
Inputs engulfed by OPCs localize to lysosomes. (A) Confocal images of an OPC from an NG2-CreER^T2^; TdTomato mouse (pseudocolored in green) immunostained for TC inputs (Vglut2, magenta) and the lysosomal marker Lamp2 (yellow). Above, volumetric reconstruction of merged signals. White arrows, points of colocalization between Vglut2 and Lamp2. Magenta arrows, Vglut2 without Lamp2. Scale bar, 5 μm. Inset scale bar, 5 μm. (B) <u>S</u>tructured <u>I</u>llumination <u>M</u>icroscopy (SIM) images and their respective reconstructions of AAV-hSYN-eGFP+ thalamocortical inputs (magenta) and NG2-CreER^T2^; TdTomato+ OPC (green) with Lamp2+ lysosomes (blue). Scale bar, 16 μm. Inset scale bar, 1 μm. (Bi) Increased magnification of Vglut2+ inputs that are not associated with Lamp2. Scale bar, 1 μm. (Bii) and (Biii) orthogonal views demonstrating presynaptic material completely internalized by the OPC. Scale bars, 1 μm. (C) Schematic of the AAV-hSYN-pSynDig virus and confocal images demonstrating the quenching of eGFP fluorescence selectively within Lamp2+ lysosomes (white arrows). Scale bar, 5 μm. (D) Quantification of the mCherry and eGFP signal at inputs outside of and within lysosomes normalized to the mCherry signal. The eGFP signal was significantly decreased compared to mCherry signal only at inputs within lysosomes. Connected points represent data from one image. Two-way ANOVA (Signal: p < 0.0001; Localization: p > 0.05; Interaction between signal and localization: p > 0.05) with Tukey post-hoc test. n = 11 images from 3 mice. (E) Confocal images of an NG2-stained OPC (green, right middle), pSynDig expressing inputs (magenta and white; right top), and an image of the pSynDig-expressing inputs within the volume of the OPC (right bottom). Scale bar, 10 μm. Left, volumetric reconstruction of OPC and pSynDig-expressing inputs. Scale bar, 10 μm. (F) Quantification of pSynDig input signal within OPCs. Ratio paired t-test mCherry versus eGFP: p < 0.05; n = 25 cells from 3 mice. (G) Reconstruction of an OPC (yellow) and pSynDig fluorescent signal (intact inputs, white; inputs being digested, magenta) in images taken on an Airyscan superresolution microscope. Lines demonstrate the location along the reconstructed OPC from which the cross-section image on the right is taken. Lines are color-matched to the borders of the cross-sections. In panels on the right, the OPC volume is outlined in yellow. Scale bar, 2 μm. Cross-section scale bar, 1 μm. *p< 0.05, ***p<0.001.

To assess the localization of internalized inputs by another strategy, we generated a fluorescent reporter of lysosomal digestion in which the presynaptic protein Synaptophysin is fused to mCherry and eGFP. Because eGFP fluorescence (but not mCherry fluorescence) is substantially quenched in acidic environments, this viral construct, which we named AAV5-hSYN-pSynDig (<u>p</u>robe for <u>Syn</u>aptic <u>Dig</u>estion), allowed us to simultaneously visualize lysosome-associated inputs (mCherry+; GFP-) and inputs outside of lysosomes (mCherry+; GFP+)(Fig. 2C,D and Extended Data Fig. 4)^17^. We observed both GFP+ and GFP-inputs within OPCs both by standard confocal and superresolution microscopy (Fig. 2E-G), confirming that a substantial number of inputs engulfed by OPCs localize to acidic compartments that are likely to represent lysosomes. In combination with our finding that presynaptic components within OPCs are smaller than those that remain unengulfed, these data suggest that OPCs engulf and digest presynaptic TC terminals in V1.

While initially focusing our analysis on synapse engulfment by OPCs in the mature brain, the observation that OPCs and microglia also engulf synapses at P20 (the beginning of the critical period of sensory-dependent refinement) and P27 (the height of sensory-dependent refinement) suggested that one or both of these cell types may contribute to the elimination of synapses in response to experience during late postnatal development. To assess this possibility, we subjected mice to a sensory deprivation and stimulation paradigm in which mice are reared in complete darkness between P20 and P27 (late-dark-rearing, LDR) then acutely re-exposed to light for ten hours (LDR+10; Fig. 3A). This is a widely used paradigm that effectively activates robust patterns of sensory-driven neural activity in the dLGN and V1^18,19^. We found that, while microglia do not change their level of engulfment as a result of sensory deprivation or stimulation (Extended Data Fig. 5), the distribution of structural contacts between microglia and OPCs shifted toward the distal ends of the OPC arbor in dark-reared mice, then returned to normal following light re-exposure (Fig. 3B,C). This result led us to hypothesize that sensory experience may coordinate signaling interactions between microglia and OPCs, some of which may be contact-dependent, to influence OPC-mediated synapse engulfment.

**Figure 3.**
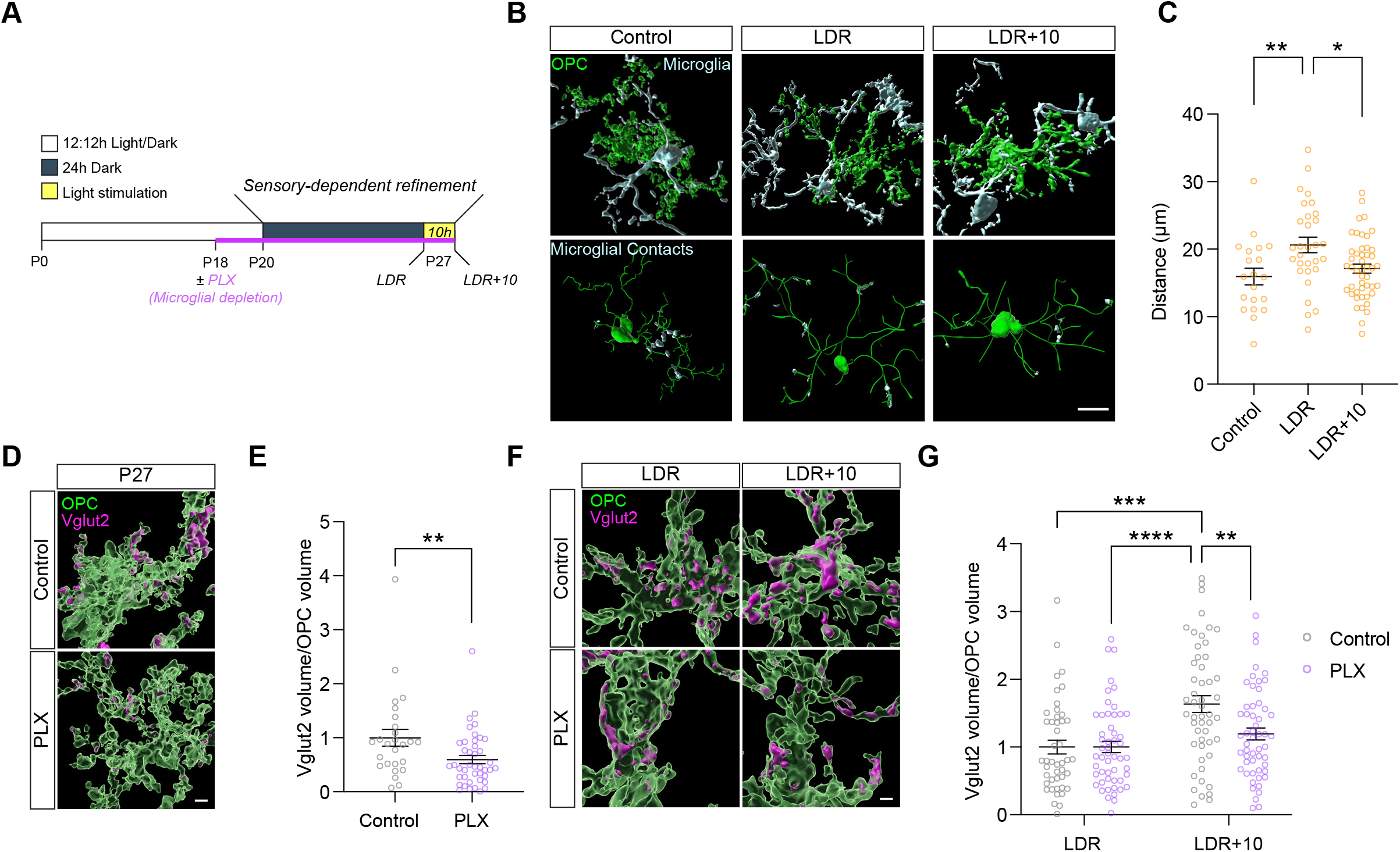
The engulfment of synaptic inputs by OPCs is regulated by sensory experience and microglia. (A) Schematic of the late dark rear (LDR) visual deprivation/stimulation and microglial depletion paradigm. (B) Top, volumetric reconstructions of OPCs stained for NG2 (green) and microglia stained for Iba1 (cyan) demonstrating physical contacts between the two cell types. Bottom, skeletonized OPC somas and processes (green) with microglial contact points shown in cyan. Control, mice reared according to a standard 12-hour light/dark cycle analyzed at P27. LDR, mice reared in darkness between P20 and P27. LDR+10, mice reared in the dark between P20 and P27 then acutely re-exposed to light for ten hours. Scale bar, 10 μm. (C) Quantification of the distance of the OPC:microglia contacts from the center of the OPC soma. One-way ANOVA (p < 0.05) with Tukey’s post-hoc test; n (OPC): P27 = 20, LDR = 30, LDR+10 = 47, from 3 mice per group. (D) Reconstructions of OPCs (green) and engulfed TC inputs (magenta) in mice at P27 following depletion of microglia between P20 and P27 via PLX5622 (or control chow) administration. Scale bar, 2 μm. (E) Quantification of the volume of synaptic material contained within each OPC in the presence or absence of microglia. Mann-Whitney t-test; p < 0.01; n (OPC): Control = 26, PLX = 42 from 3 mice per group. (F) Reconstruction of OPCs (green) and synaptic inputs (magenta) as shown in (D) from dark-reared (LDR) and visually stimulated (LDR+10) mice. Scale bar, 2 μm. (G) Quantification of synaptic engulfment as shown in (E) in LDR and LDR+10 mice containing or lacking microglia. Two-Way ANOVA (Stimulation: p < 0.0001; Microglia: p < 0.05; Interaction between stimulation and microglia: p < 0.05); n (OPCs, Control/PLX): LDR = 45/52, LDR+10 = 51/54 from 3 mice per group. (C), (E), and (G), Bars represent mean +/− SEM. *p< 0.05, **p< 0.01, ***p<0.001, ****p<0.0001.

To test this hypothesis, we took advantage of PLX5622, an inhibitor of Colony Stimulating Factor Receptor 1 that, when administered to mice, depletes microglia from the brain almost entirely within three days of beginning administration (Extended Data Fig. 6). We found that depletion of microglia in normally reared mice between P20 and P27 significantly decreased the amount of synaptic material found within OPCs (Fig. 3D,E). Furthermore, whereas microglial engulfment was not altered by changes in sensory experience, synapse engulfment by OPCs in control mice was significantly heightened following light re-exposure after dark-rearing. Remarkably, this experience dependent increase in OPC-mediated engulfment was dampened in the absence of microglia, indicating that sensory experience and microglia converge to promote the OPC-mediated engulfment of TC inputs (Fig. 3F,G). In addition to an overall increase in engulfment, analysis of the distribution of discrete sites of engulfment across the highly complex OPC arbor revealed a sensory-driven shift in engulfment sites toward the more distal regions of OPC processes. This shift occurred in response to either sensory deprivation or stimulation, suggesting that bidirectional manipulations of experience may position OPCs to eliminate synapses across a broader cortical volume (Extended Data Fig. 7). Altogether, these data demonstrate that OPCs engulf synaptic inputs in response to sensory experience during a critical period of sensory-dependent plasticity, and that microglia provide signals to OPCs to promote their engulfment of synapses in an experience-dependent manner.

Despite a growing number of studies describing the roles of glia in synaptic refinement in the early postnatal brain, how non-neuronal cells remodel synaptic connectivity during later experience-dependent phases of development and in the adult remains an area of active investigation^20,21^. Here, we uncover a role for OPCs in shaping synaptic connectivity by engulfing synaptic inputs in sensory cortex in the late postnatal and mature mammalian brain. Our findings are consistent with the observation of axonal structures within OPCs as assessed by electron microscopy, and the regulation of axonal remodeling by OPCs in the zebrafish tectum^22,23^.

Here, we place these observations within an important neurobiological context by showing that OPCs engulf more synaptic material when visual experience is heightened, and that this sensory-dependent increase in OPC-mediated engulfment requires microglia. The ability of OPCs to detect changes in sensory experience is consistent with previous reports demonstrating that sensory input, and neuronal activity more generally, regulate OPC maturation and proliferation, as well as oligodendrogenesis^24,27^. Additionally, microglia are known to contribute to OPC survival, development, and maintenance in adulthood^28,29^. In combination with these studies, our data suggest that experience can impact OPCs in multiple ways, not only by driving adaptive myelination but also by triggering OPCs to directly remodel synaptic connectivity through the engulfment of presynaptic terminals. Given mounting evidence that deficits in the functions of oligodendrocyte lineage cells exacerbate neurological disorders associated with synapse loss including Alzheimer’s disease and multiple sclerosis^30,31^, our discovery that OPCs can refine synapses through the engulfment of presynaptic inputs is likely to shed light on mechanisms of disease in the human brain.

## Acknowledgements

We thank Drs. Ullas Pedmale and Daniele Rosado for the use of their Airyscan microscope, funded by NIH grants R35GM125003, GM12500303S1, and GM12500304S1. In addition, we thank Drs. Beth Stevens, Cody Walters, and Ullas Pedmale for critical feedback on the manuscript. Erika Wee, director of the microscopy core at Cold Spring Harbor Laboratory, assisted with confocal and SIM microscopy imaging. This work was funded by a Rita Allen Scholar Award, a McKnight Scholar Award, and a Klingenstein-Simons Fellowship to L.C.

## Author contributions

L.C. conceptualized the study. Y.A. generated the initial discovery that OPCs engulf synaptic inputs, and was instrumental in overseeing experiments performed by other authors. A.F. performed and analyzed two-photon microscopy experiments. J.D., U.V., J.K., and A-S. N. generated and analyzed experimental data. L.C. wrote the first draft of the manuscript after which all authors contributed to its editing.

## Methods

### Animal Models

All experiments were performed in compliance with protocols approved by the Institutional Animal Care and Use Committee (IACUC) at Cold Spring Harbor Laboratory according to protocol #20-3. The following mouse lines were obtained from the Jackson Laboratory: C57Bl/6J (JAX: 000664), NG2-CreER^T2^ (Cspg4-Cre, JAX: 008538), and Rosa26-CAG-LsL-TdTomato (JAX: 007914; Ai14). NG2-CreER^T2^ mice were bred with Ai14 mice in-house to yield NG2-CreER^T2^; TdTomato mice in which oligodendrocyte precursor cells (OPCs) are labeled with TdTomato upon tamoxifen (TAM) administration. Except when noted, animals were housed in normal 12:12 hour light-dark cycles. Analyses were performed on equal numbers of male and female mice at postnatal days (P)10, P20, P27, and P90. Live imaging was performed on animals between 2 and 6 months of age. No sex differences were observed in this study.

### Sensory Deprivation and Stimulation Paradigm

To study the effects of sensory experience on synapse engulfment by OPCs and microglia, C57Bl/6J mice were reared according to a standard 12-hour light/dark cycle until P20, at which point they were weaned from their mothers and separated into experimental groups. One cohort continued to be housed in normal light conditions until P27. Two cohorts of mice were placed in a well-ventilated light-proof cabinet (Actimetrics) until P27 (late-dark-reared, LDR). One of the two cohorts subjected to LDR was harvested at P27 in the dark by an investigator using night vision goggles while the other was acutely re-exposed to light for ten hours, then harvested. These cohorts are referred to as LDR and LDR+10 throughout the paper.

### Plexxicon 5622 Administration

To study the role of microglia on OPC function, mice were fed irradiated 1200 mg/kg freebase Plexxicon 5622-formulated chow (PLX; Research Diets, Inc.: D1110404i), which blocks Colony Stimulating Factor Receptor 1, or control chow produced in parallel (D19101002) from P18 to P27 to pharmacologically deplete microglia from the brain. Mice were fed on the chow ad libitum and the investigator provided all husbandry for the mice during treatment.

### Tamoxifen Administration

Tamoxifen (Sigma: T5648) was dissolved in sunflower oil (Sigma: S5007) overnight at 37 °C with shaking to achieve a working concentration of 20 mg/mL. Animals were administered a bolus of TAM at 150 mg/kg on two consecutive days at a minimum of two weeks before imaging and analysis.

### Immunofluorescence

Animals were anesthetized with isoflurane and perfused with ice cold Phosphate Buffered Saline (PBS) followed by 4 % paraformaldehyde (PFA). Brains were removed and incubated in 4% PFA overnight at 4° C. The next day, the brains were washed by rotating in fresh PBS 3 times for 10 mins and then allowed to sink in 15% and then 30% sucrose in PBS at 4° C. In preparation for sectioning, brains were embedded in OCT (VWR: 25608-930) and stored at −80° C. 25 μm thick coronal or sagittal sections containing the primary visual cortex and dLGN were collected onto Superfrost Plus microscope slides (Thermo Fisher Scientific: 1255015) using a cryostat and stored at −80° C.

For staining, the sections were washed at room temperature (RT) for 10 mins in PBS, and then dried for 10 mins at 60° C. A hydrophobic barrier was drawn around the samples using an ImmEdge Pen (VWR: 101098-065). A blocking solution (5 % Fetal Bovine Serum [FBS] or Normal Goat Serum [NGS] and 0.3% Triton-X100, in PBS) was applied for 1 hour at RT. Next, the blocking solution was replaced with primary antibodies prepared in a probing solution (5% FBS or NGS and 0.1% Triton in PBS). Primary antibodies were incubated at 4° C overnight (O/N) in most cases, and for 48 hours for staining with rat α NG2 (1:250; Thermo Fisher Scientific: MA5-24247). Other primary antibodies used include guinea pig α Vglut2 at 1:1000 (Sigma: AB2251-l); rabbit α Iba1 at 1:1000 (Wako: 019-19741); rat α Lamp2 at 1:200 (Abcam: AB13524); and rabbit α Sox10 at 1:100 (Abcam: AB227680). After primary incubation, the tissue was washed 4 times with washing solution (PBS adjusted to 0.1% Triton) for 10 mins per wash. The following Alexafluor secondary antibodies (Invitrogen) were diluted to 1:1000 in probing solution and incubated on the tissue for 1 hour at RT: goat α guinea pig 647 (A21450); goat α guinea pig 555 (A21435); donkey α rabbit 405 (A21450); goat α rabbit 488 (A11008); goat α rabbit 647 (A21428); goat α rat 405 (A48261); donkey α rat 488 (A21208); and goat α rat 647 (A21247). Following secondary incubation, the sections were washed 4 times with washing solution, mounted with Flouromount-G (SouthernBiotech: 0100-01), and coverslipped.

### Confocal Imaging Parameters

In most cases, confocal images were acquired on an LSM710 or LSM780 (Zeiss) microscope with 20x (air) and 63x (oil) objectives. For engulfment analyses, the microscope was centered over layer 4 of visual cortex (V1) at 20x, identified by a Vglut2+ band, before transitioning to 63x. Z-stack images were acquired to capture several OPCs and/or microglia. To highlight the expression of phagocytic genes in OPCs, *Pdgfra+* nuclei were imaged at 63x.

### NG2-Cre^T2^;TdTomato Validation

Fixed brain sections from NG2-Cre^T2^; TdTomato mice were stained for Sox10 and NG2 as described above, and imaged on a confocal at 20x. To determine the percentage of TdTomato+ cells that were either within the oligodendrocyte lineage (Sox10+) or putative OPCs (NG2+), maximum projections of the images were generated to be manually counted with the “Cell Counter” plugin in ImageJ. First, markers were placed at all TdTomato+ cells. Next, at each TdTomato+ marker location, the investigator determined and tabulated whether the location was also Sox10+ or NG2+.

### Fluorescence *In Situ* Hybridization (FISH)

Brains were fixed by perfusion in 4% PFA, embedded in OCT and stored at −80° C until processing. Sections of 25 μm thickness were mounted on Superfrost Plus slides. Multiplexed single-molecule FISH was performed using the RNAscope platform V2 kit (Advanced Cell Diagnostics [ACD]: 323100) according to the manufacturer’s protocol for fixed-frozen sections. The samples were mounted with Flouromount-G with DAPI (SouthernBiotech: 0100-20), and coverslipped. Commercial probes obtained from ACD detected the following genes: *Pdgfra* (480661-C2), *Mertk* (441241-C3), *Calcrl* (452281-C3), *Arsb* (837631), and *Ptprj* (883051).

### Re-analysis of Single-cell RNA-sequencing Data

Data from Hrvatin et al, *Nature Neuroscience,* 2018 were downloaded as a raw counts matrix from the Gene Expression Omnibus database (GSE 102827). While this dataset includes cells from mice that were dark-reared, dark-reared then re-exposed to light for 1 hour, and dark-reared then re-exposed to light for 4 hours, none of the genes we plotted were differentially expressed between conditions so we plotted their expression across all cells in the dataset. The data were processed via the Seurat v3 pipeline using standard parameters. OPC clusters were identified by enriched expression of *Pdgfra*. For Figure S2, transcripts were plotted across the UMAP using the FeaturePlot function in Seurat.

### Engulfment Quantification

Preprocessing of 63x images stained for OPCs and/or microglia was performed in ImageJ. First, the “Enhance Contrast” command was run so that 0.1% of pixels would be saturated, and then a mean filter with pixel radius of 1.5 μm was applied. Next, a region of interest (ROI) was drawn around a given cell, cropped into its own file and used in downstream Imaris (BitPlane) processing and analysis. In Imaris, volumetric reconstructions of the fluorescence images were created using both the “Spots” and/or the “Surfaces” Objects. A Surface Object was used to reconstruct a cell of interest following the guided creation wizard. The investigator then deleted any discontinuous part of the Surface that could not be clearly traced back to the soma of the cell, using the fluorescence as reference. Next, this cell Surface was used as the ROI to create a mask of target channels (e.g. Vglut2, Lamp2), defined by the signal included within the Surface. New Surface Objects were generated using these masks, which represent the internal contents of the cell. Across conditions within a biological replicate the same creation parameters were used in generating the internalized Surfaces. The volumes of the cell and internalized Surfaces were collected from the statistics tab in Imaris. The internalized volume was then normalized to a cell’s volume to represent the amount of engulfment by a given cell.

In some cases, masks of the Lamp2 channel within an OPC Surface was made to reconstruct lysosomes. To quantify lysosome contents, the same internalization approach was used where the lysosomal Surface was treated as the “cell” to mask the target channel.

To quantify the distance of engulfment loci from the center of an OPC, we first defined the center of the cell by manually placing a Spots Object at the soma of the reconstructed cell using the auto-depth-based-on-fluorescence option. Next, a Spots Object of the masked Vglut2 signal was created using the wizard, with a Spots-diameter of 2 μm. This approach did not place spots at every Vglut2 puncta, but instead placed a Spot near collections of Vglut2 puncta. The predefined “Shortest Distance to Spots” statistic was used to measure the distances between the engulfment loci and the center of the cell. To quantify the range of synaptic surveillance by a given OPC, the axis-aligned bounding-box statistic of the engulfment loci Spots was collected.

### Structured Illumination Microscopy (SIM)

Brains were fixed in 4% PFA then embedded in OCT and stored at −80° C. Sections of 25 μm thickness were subjected to immunofluorescence in a free-floating format. Each section was placed in one well of a 24-well plate. Sections were stained with primary and secondary antibody as described under “Immunofluorescence” above, except in larger volumes of 1 mL. Stained sections were mounted onto thickness no. 1 ½ High-performance Zeiss cover glasses (Thermo Fisher Scientific: 10474379) and then centered onto Superfrost Plus microscope slides with ProLong™ Gold Antifade Mountant (Thermo Fisher Scientific: P36934). Stained samples were kept at 4° C until imaging. 3-D structured illumination microscopy (3-D SIM) of fixed, stained samples were acquired using an Applied Precision V3 OMX system equipped with a 100x/1.4 NA U-PLANAPO objective (Olympus) and two Cascade II^®^ 512 EM-CCD cameras (Photometrics). Stacks of 6 optical sections (125 nm step) were acquired consecutively in two channels (488 nm and 593 nm) using DeltaVision software (Applied Precision). 3-D super-resolution image stacks were reconstructed using SoftWorx 6.5.2 using channel specific OTFs and Wiener filter settings of 0.001 or 0.002. These image stacks were then imported into Imaris for volumetric reconstructions.

### Airyscan Imaging

Samples were imaged on a Zeiss LSM900 microscope using the Airyscan mode with a 63x objective. The image underwent Airyscan Processing with Auto Filter selected within Zeiss Zen. The processed image was then transferred to Imaris for volumetric reconstruction as described above.

### pAAV:hSYN-Synaptophysin-mCherry-eGFP (pSynDig)

We purchased the reporter of presynaptic ATP Syn-ATP (Addgene: 51819) and the fluorescent marker hSYN-eGFP (Addgene: 50465) plasmids for downstream creation of the <u>p</u>robe for <u>Syn</u>aptic <u>Dig</u>estion (pSynDig) construct. We amplified an elongated coding region for mCherry from Syn-ATP (forward primer gcgcagtcgagaaggtaccgGCAGCAATGGACGTGGTG; reverse primer: ccttgctcaccatggtggcgGGTCCCTTGTACAGCTCG). The elongated coding region (1728 bp) was then gel purified using the Qiagen MinElute Gel Extraction Kit (28604) and subjected to Gibson assembly via the New England Biosciences HiFi DNA Assembly Cloning Kit (NEB: E5520S) to insert the amplified region into the hSyn-eGFP vector following its linearization with BamHI (NEB: R0136S). The resulting plasmid was gel-purified and sequenced before packaging into an adeno-associated virus (AAV) to yield AAV5-hSYN-pSynDig at a titre of 1.2 x 10^14^ gc/mL.

### pSynDig Validation and Quantification

We confirmed that pSynDig injections into the dLGN labeled Vglut2+ thalamocortical inputs in layer 4 of visual cortex (V1) by imaging mCherry, eGFP, and Vglut2+ puncta (following immunostaining with an antibody against Vglut2) in confocal images taken with a 63x objective at Nyquist settings for downstream deconvolution using Huygens Essentials (SVI). Images were deconvolved based upon imaging parameters (e.g. pixel resolution, excitation wavelength, number of excitation photons, depth of acquisition, numerical aperture of objective and medium, and emission wavelength) and the Huygens express deconvolution wizard set to conservative deconvolution as means of increasing image resolution and signal to noise. We measured the intensity of each channel using line intensity quantification in ImageJ. We observed largely overlapping mCherry and eGFP signal colocalized with Vglut2, as expected. To further validate this, we used the Imaris Coloc function to measure the degree of colocalization of both the eGFP and mCherry signals using the mean Pearson’s correlation coefficients between 3 images per animal. As expected, we observed a small percentage of puncta that were mCherry+ but eGFP-. To verify that these mCherry+, eGFP-puncta represented inputs in the process of lysosomal degradation, we immunostained the tissue for the lysosomal marker Lamp2 and took 3-dimensional images (z-stacks) on a confocal microscope using a 63x objective. In post-processing, we applied a gaussian blur of 0.132 μm. Maximum projections of the z-stacks were made, then the mean intensities of the mCherry, eGFP, and Lamp2 channels were quantified. Images were moved into Imaris, where lysosomes were reconstructed using a Surface Object. In the wizard, the recorded mean intensities from ImageJ were used as the threshold value to define the Surfaces. Masks of the eGFP and mCherry channels were made from the lysosome Surfaces to analyze the within-lysosome pSynDig signal. Surfaces of the masked eGFP and mCherry signals were created, this time using a 0.75*(recorded-mean-intensity) as the threshold. To quantify the intensity of mCherry and eGFP outside of lysosomes, the masked channels were subtracted from the original channels, thereby removing any signal that was within a lysosome. mCherry and eGFP Surfaces were generated from these subtracted channels, using the same wizard thresholds as for the within-lysosome group. The sum intensity statistic of the eGFP and mCherry signals were collected for the within- and outside-lysosome Surfaces, before being normalized to their respective mCherry signal. The example image in figure 2C was taken with a 63x objective with 1.5 x digital zoom, before being deconvolved for clarity.

To quantify pSynDig in OPCs, the same approach was used as for the within-lysosome group, where OPC Surfaces were reconstructed and used to mask the mCherry and eGFP channels. In figures 2E and F, only mCherry Surfaces are shown and were pseudo-colored white or magenta if they overlapped with an eGFP Surface or not, respectively.

### OPC:Microglia Contact Quantification

The same ImageJ pre-processing as above was applied to images, centered on an OPC then cropped to include the OPC and surrounding microglia. Surfaces for both cell types were created. OPC Surfaces were manually filtered to remove objects that did not originate from the soma. For the microglial signal, however, surfaces were kept even if the soma was not visible to avoid omitting important interactions between the processes of the two cell types. From the OPC Surface, the “Surface-Surface Contact Area” Imaris XTension (https://github.com/Ironhorse1618/Python3.7-Imaris-XTensions/blob/master/XT_MJG_Surface_Surface_ContactArea2.py) was run. Briefly, it creates a 1-voxel-thick shell of the OPC and the microglia, then masks where the two intersect. From this mask, a 1 voxel-thick Surface is created of the contact area. For consistency, we quantified the interaction with the volume statistic of the Surface instead of the surface area to avoid double counting each side of the Surface and adding the superfluous area created by the end-caps of the Surface. The volume was then normalized to the volume of the OPC to account for variations in OPC size. The skeletal representations of the OPCs were constructed using the Filaments Object for figure purposes only (Fig. 3B).

### AAV injections

Mice aged between 2-6 months were injected with a slow-release analgesic, meloxicam [2.5 mg/kg, subcutaneous] before being anesthetized using isoflurane (SomnoSuite, Kent Scientific; 3-5% induction, 1-2% maintenance). Once anesthetic depth was achieved, mice were placed onto a stereotaxic apparatus where body temperature was maintained using a heating pad. Mice were then unilaterally injected with either AAV9-hSYN-eGFP (Addgene viral prep # 50465-AAV9) or AAV5-hSYN-pSynDig; 500 μL with a flow rate of 50 nL/min) into right hemisphere dLGN (2.15x, −2.15y, −2.9z mm from bregma). Following surgery, animals were administered Flunixin (10 mg/kg) and allowed to recover on a heating pad before returning them to their home cages.

### Chronic Window Implantation

Mice (n = 10) aged between 2-6 months, previously injected with meloxicam [2.5 mg/kg, s.q.] were anesthetized using isoflurane (3-5% induction, 1-2% maintenance) and body temperature was maintained with a heating pad throughout surgery and during initial recovery. After initial AAV injection (as described above), a >3 mm diameter craniotomy was drilled using a dental drill (RWD: 78001) over primary visual cortex approximately +2.5 mm lateral, and −2.9 mm posterior from Bregma. A 3 mm glass coverslip, sterilized with 70% ethanol, was then placed over the craniotomy and a mixture of surgical glue (Vetbond: 3M) and cyanoacrylate glue was used to secure the coverslip onto the skull. The skull was covered with a thin layer of Vetbond and then sealed with dental cement (Ortho-Jet: Land Dental). Finally, a custom-made head bar was secured onto the skull using luting cement (Metabond: C&B). Mice were then administered flunixin meglumine [10mg/kg, i.p.], and allowed to recover on a heating pad until ambulatory and then were allowed to recover for 1-2 weeks before imaging. After recovery, the quality of the windows was checked before imaging, and mice with suboptimal windows were euthanized and used for downstream immunofluorescence quantification in fixed tissue.

### *In Vivo* 2-photon Imaging

After recovery from window implantation (1-2 weeks), mice were secured into a custom head mount and movement restraint system. Mice were imaged using a custom 2-photon system (Independent Neuroscience Services) with a ThorLabs tunable Tiberius laser. Laser wavelength was tuned to 980 nm to image both TdTomato and eGFP concurrently. We utilized a 16x (N.A. 0.8) water immersion lens (Nikon), and light was captured using two photomultiplier tubes fitted with filters (520-565, and >565 nm) for eGFP and TdTomato respectively. While imaging between (150 and 350 μm depth from the pia) the laser power was kept below 30 mW to avoid photo-damage. Imaging volumes were captured at near Nyquist settings (either 512×512, and 1024×1024 for time-lapse and single time point recordings respectively; resulting in voxels ≤ 264 × 264 × 1000 nm) were selected for fields with TdTomato+ cells with distinguishable OPC morphology and thalamocortical inputs. For time-lapse recording, volumes were taken once per minute. Raw image files were then processed using Huygen’s software (SVI, Netherlands) for crosstalk correction, registration along the z and t dimensions, and then processed with their multiphoton deconvolution wizard (set to conservative deconvolution as previously done for pSynDig experiments). Crosstalk-corrected, registered and deconvolved images were then imported into Imaris and Surface Objects of OPCs and thalamocortical inputs were created, with surfaces of OPCs being limited to the OPC soma and major process. Thalamocortical inputs were then classified by their distance to the OPC’s surface using the “Shortest Distance to Surfaces” filter function in Imaris, with thalamocortical inputs that were greater than 0 nm classified as non-contacting inputs, inputs with a distance to OPCs 0 nm classified as contacting inputs, and inputs with a distance of less than −270 nm classified as “engulfed” inputs. The classified thalamocortical inputs’ volume were then averaged per video and per time frame, and then normalized to the average volume of non-contacting inputs over the entire imaging period, where applicable. For data presentation, the OPC surface volume was pseudocolored with green and classified input volumes were pseudocolored using the, “Overlapped Volume Ratio to Surface” function.

#### Note on identifying OPCs in vivo

Because we typically imaged NG2-CreER^T2^; TdTomato mice following greater than 2 weeks post-TAM, some of the Cre-expressing OPCs had differentiated to label mature oligodendrocytes with TdTomato by the time of imaging. However, OPCs could be easily distinguished from oligodendrocytes based upon their oblong, bean-shaped somata compared to oligodendrocytes, the somata of which were more spherical (see Extended Data Figure 1).

### Blinding

Experimenters were blinded to conditions for quantitative imaging experiments. One experimenter harvested the tissue and assigned it a randomized label before providing the blinded tissue to another experimenter for analysis. After data acquisition and processing, the data were plotted in Graphpad by L.C., A.F., or Y.A. after which the samples were unblinded.

### Statistics

All statistics were performed in Graphpad by L.C., A.F., or Y.A. and are described in the figure legends.

---

**Extended Data Figure 1.**
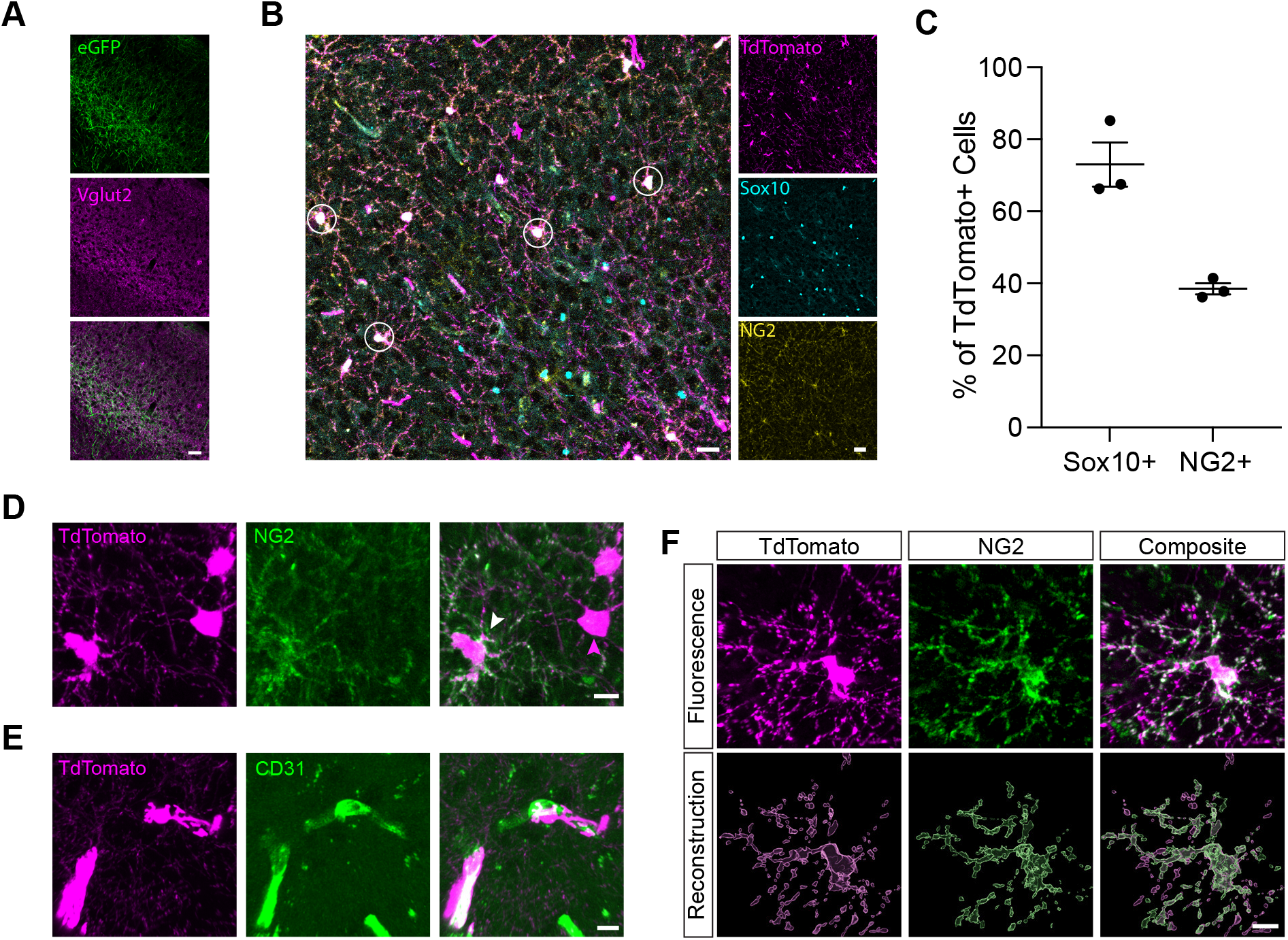
Validation of the OPC reporter line. (A) Confocal images of TC axons and inputs in V1 labeled with AAV-hSYN-eGFP (green) and immunostained for Vglut2 (magenta). Scale bar, 40 μm. (B) Representative image of TdTomato+ cells (magenta) in the NG2-CreER^T2^; TdTomato mouse line. Cells of the oligodendrocyte lineage immunostained for Sox10 (green) and the OPC marker NG2 (yellow). White circles, OPCs confirmed by co-expression of TdTomato and NG2. Scale bar, 20 μm. Scale bar fluorescence, 40 μm. (C) Quantification of the percentages of TdTomato+ cells that co-stain for Sox10 and NG2. Individual data points and mean +/− SEM shown; n = 3 mice per group. (D) Higher resolution images of TdTomato+ cells (magenta) immunostained for NG2 (green). White arrow, confirmed OPC in which TdTomato and NG2 signal overlap. Compared to other Sox10+ cells, OPCs can be distinguished by their bean-shaped somata. Scale bar, 10 μm. (E) Confocal images of rare TdTomato+ cells (magenta) co-localizing with blood vessels (CD31, green). These are likely pericytes and are easy to distinguish from OPCs based upon morphology. Scale bar, 10 μm. (F) High magnification image of a TdTomato+ OPC (magenta) immunostained for NG2 (green). Bottom, volumetric reconstructions of the same cell based upon TdTomato versus NG2 signal, demonstrating a high level of overlap between the two OPC markers. Scale bar, 10 μm.

**Extended Data Figure 2.**
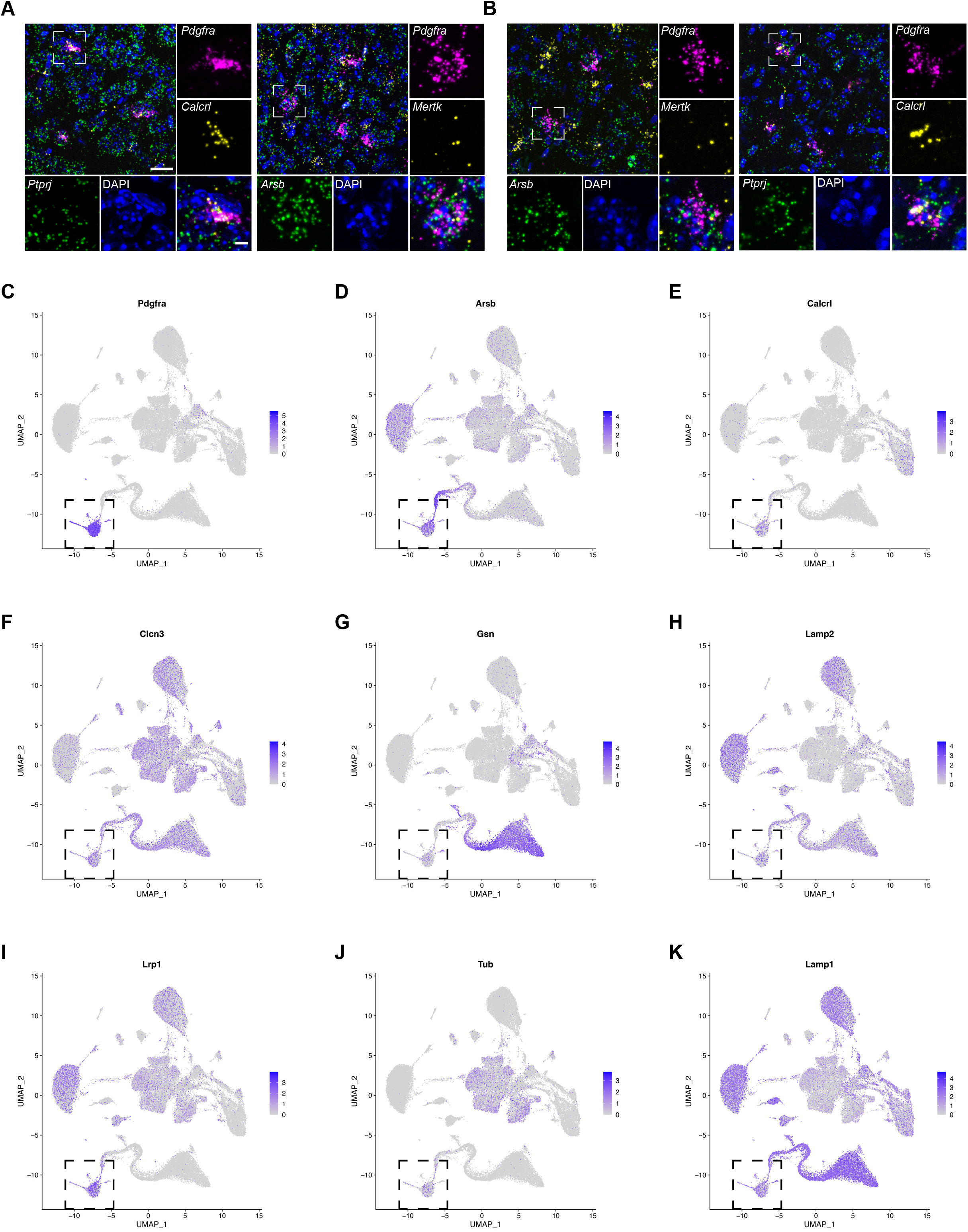
Expression of engulfment-related genes in OPCs. Confocal images of V1 sections subjected to fluorescence *in situ* hybridization (RNAscope) and probed for the OPC marker gene *Pdgfra* (magenta) along with genes that encode known regulators of phagocytic engulfment and lysosomal function: *Ptprj* (green), *Calcrl* (yellow), *Arsb* (green), and *Mertk* (yellow). DAPI shown in blue. (A) P27; (B) P90. Scale bar, 20 μm. Inset scale bar, 5 μm. (C) – (K) UMAPs demonstrating engulfment-related gene expression (taken from PANTHER gene ontology term list, “phagocytosis”) across all clusters in dataset from Hrvatin, *et al,* 2018. Gene name given above graph. Square, OPC cluster.

**Extended Data Figure 3.**
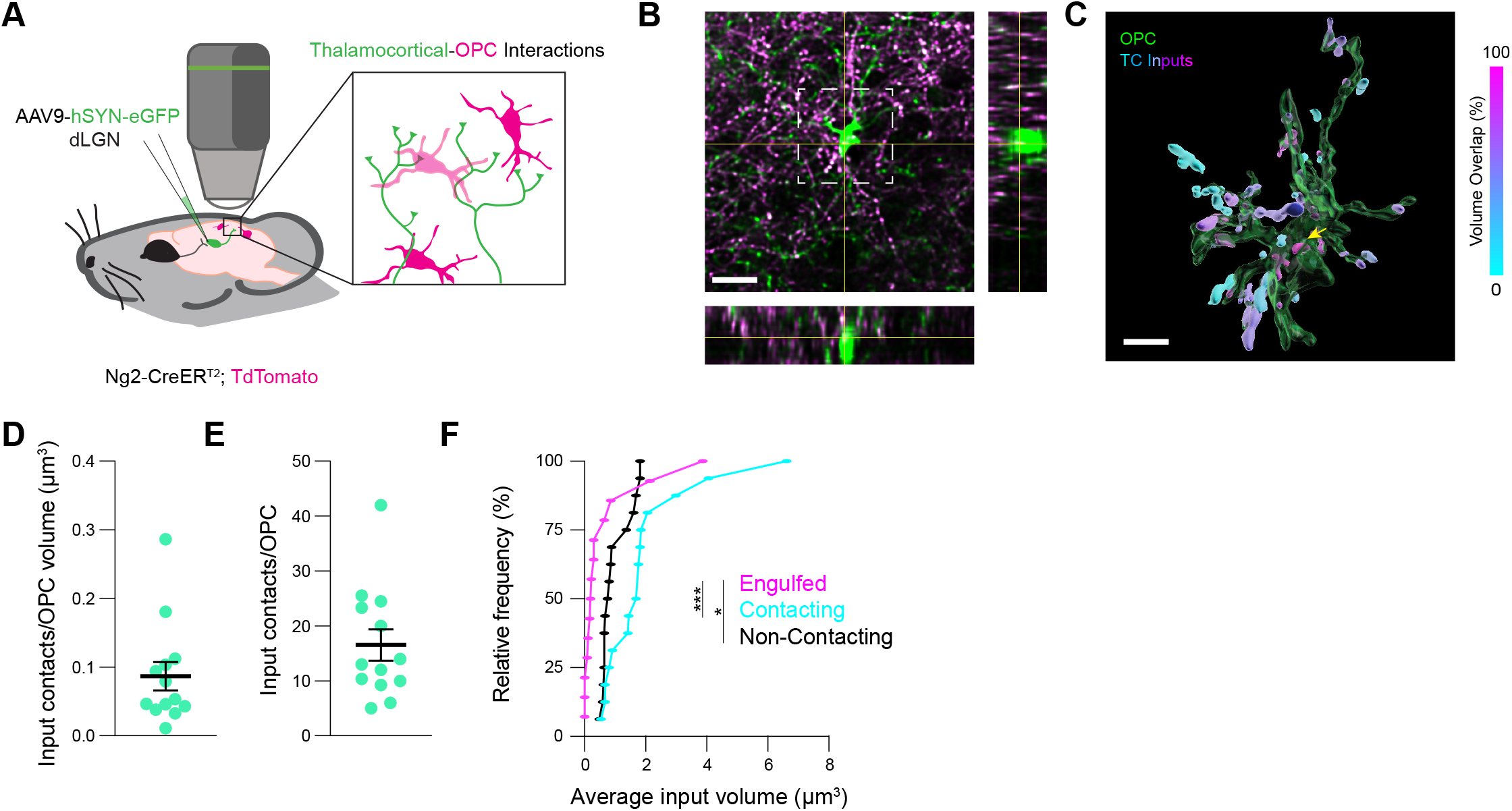
*In vivo* two-photon imaging of OPC-synapse interactions. (A) Schematic demonstrating the viral labeling of thalamocortical (TC) axons with AAV9-hSYN-eGFP in NG2-CreER^T2^; TdTomato mice and the live-imaging paradigm. (B) Maximum projection of a Z-stack of V1 in an awake mouse taken on a two-photon microscope (reconstructed in Fig. 1I). OPC, green. Inputs, magenta. Orthogonal projections on bottom and to the right. Scale bar, 20 μm. (C) Volumetric reconstruction of the OPC from (B) (green) with inputs colored based upon their overlap with OPC signal. Yellow arrow, completely internalized input. Scale bar, 10 μm. (D) The number of TC inputs contacting an OPC normalized to OPC volume. (E) The number of TC inputs contacting an OPC. (F) Cumulative frequencies of average input size categorized by contact or engulfment by OPCs. Kruskal-Wallis test with Dunn’s post-hoc comparisons, ***p < 0.001, *p < 0.05. For all data n = 15-17 volumes from 5 mice.

**Extended Data Figure 4.**
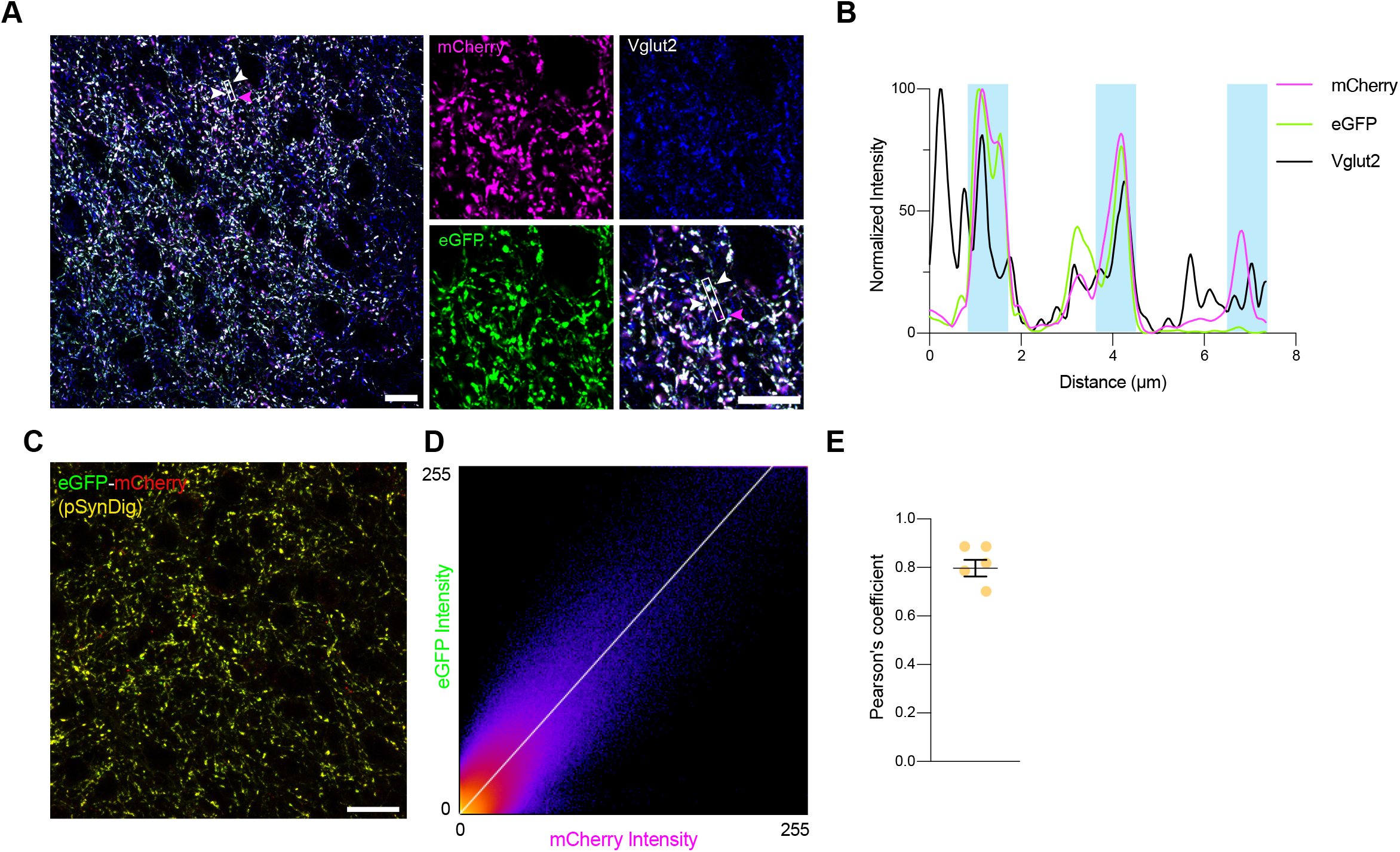
Validation of pSynDig as a marker of synaptic inputs. (A) Confocal image of layer 4 of V1 following viral infection of the dorsal lateral geniculate nucleus (dLGN) with AAV5-hSYN-pSynDig. Most pSynDig+ inputs show tight colocalization between mCherry (magenta) and eGFP (green) signal overlapping with the input marker Vglut2 (blue). White box, example region quantified in (B). White arrows, inputs that are positive for both mCherry and eGFP. Magenta arrow, input that is only positive for mCherry. Scale bar, 10 μm. Inset scale bar, 10 μm. (B) Quantification of fluorescence intensity across the line scan denoted by the white box in (A) for each channel separately, normalized to each channel’s respective maximum intensity. Note the high degree of overlap between mCherry and eGFP signal in the first two blue bars and mCherry alone in the last bar. Also note the presence of a Vglut2+ synapse represented by the black peak that precedes the first blue bar, which is likely derived from a dLGN neuron that was not infected with the pSynDig virus. (C) Confocal image of pSynDig+ inputs in V1. Scale bar, 20 μm. (D) Quantification of fluorescence intensities of eGFP and mCherry across the imaging frame shows a high degree of colocalization. (E) Quantification of Pearson’s coefficient describing the colocalization of mCherry and eGFP signal at inputs expressing pSynDig. Individual data points with mean +/− SEM. n = 5 images/3 mice.

**Extended Data Figure 5.**
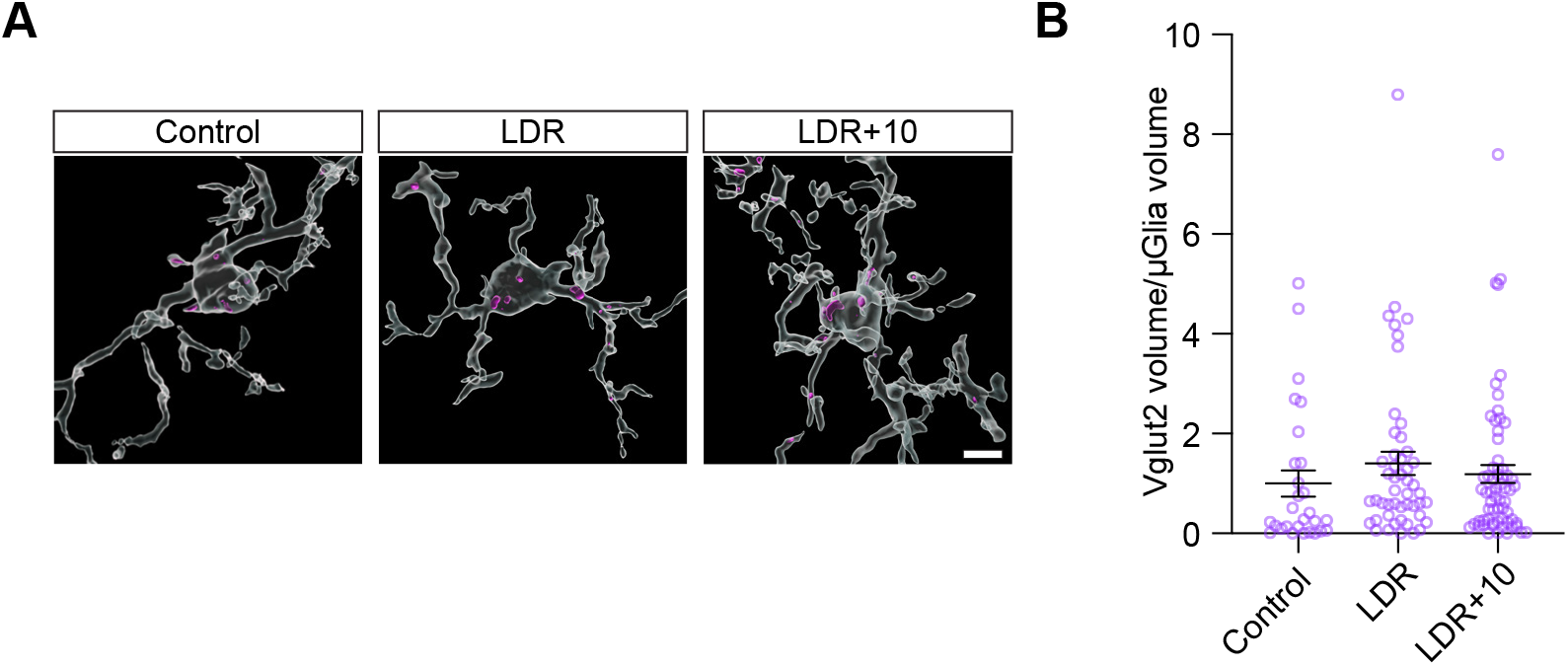
Synapse engulfment by microglia is not sensitive to robust changes in sensory experience. (A) Volumetric reconstructions of microglia immunostained for Iba1 (white) and engulfed Vglut2+ inputs (magenta) in normally reared mice at P27, mice dark-reared between P20 and P27 (LDR), and mice re-exposed to light for 10 hours following LDR (LDR+10). Scale bar, 2 μm. (B) Quantification of the volume of synaptic material within microglia of each condition. One-way ANOVA (p > 0.05) with Tukey’s post-hoc test; n (microglia): P27 = 28, LDR = 49, LDR+10 = 64, from 3 mice per group.

**Extended Data Figure 6.**
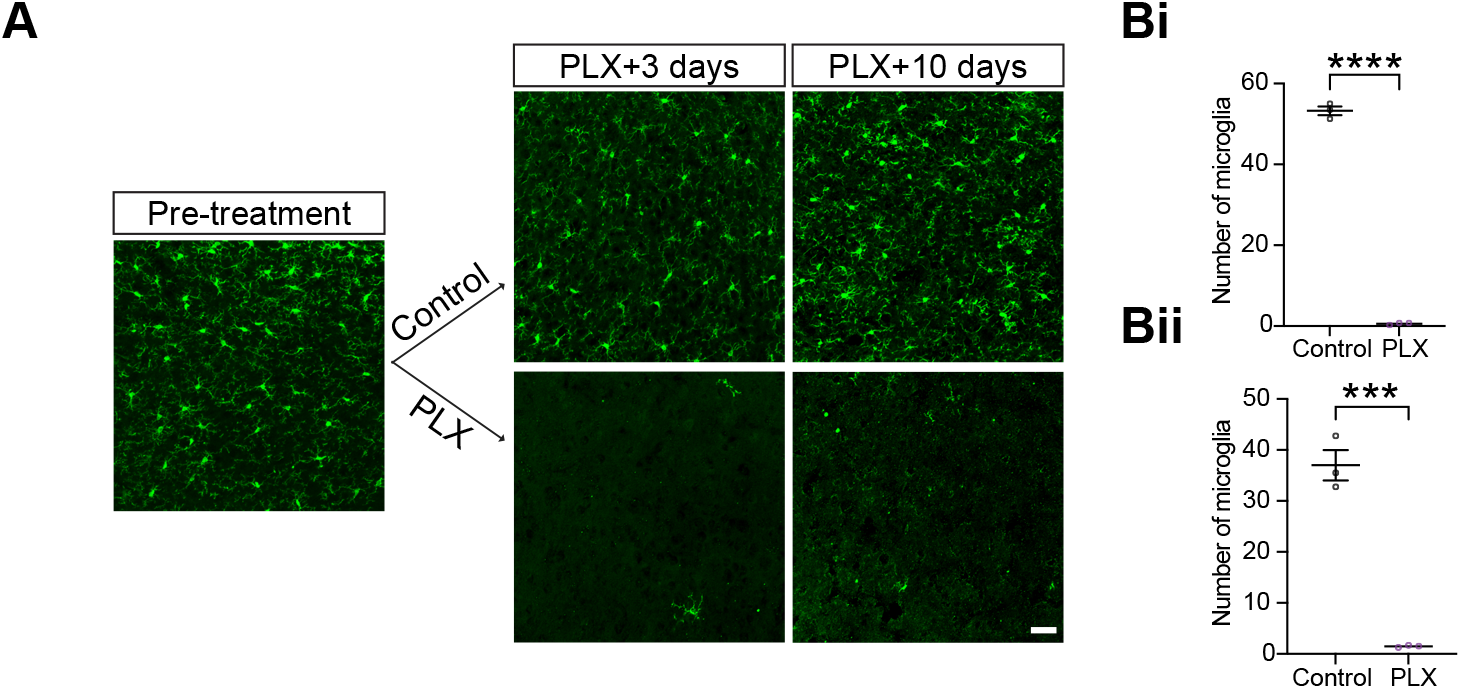
Validation of microglial depletion using PLX5622. (A) Representative confocal images of microglia immunostained for Iba1 (green) in the visual cortex before and during PLX5622 administration. Scale bar, 20 μm. (B) Quantification of the number of microglia in V1 averaged across three mice per condition. (Bi) Three days of PLX5622 administration. (Bii) Ten days of PLX5622 administration. Individual data points with mean +/− SEM. Unpaired t-test; three days, ****p < 0.0001; ten days, ***p < 0.001. n = 3 mice per group.

**Extended Data Figure 7.**
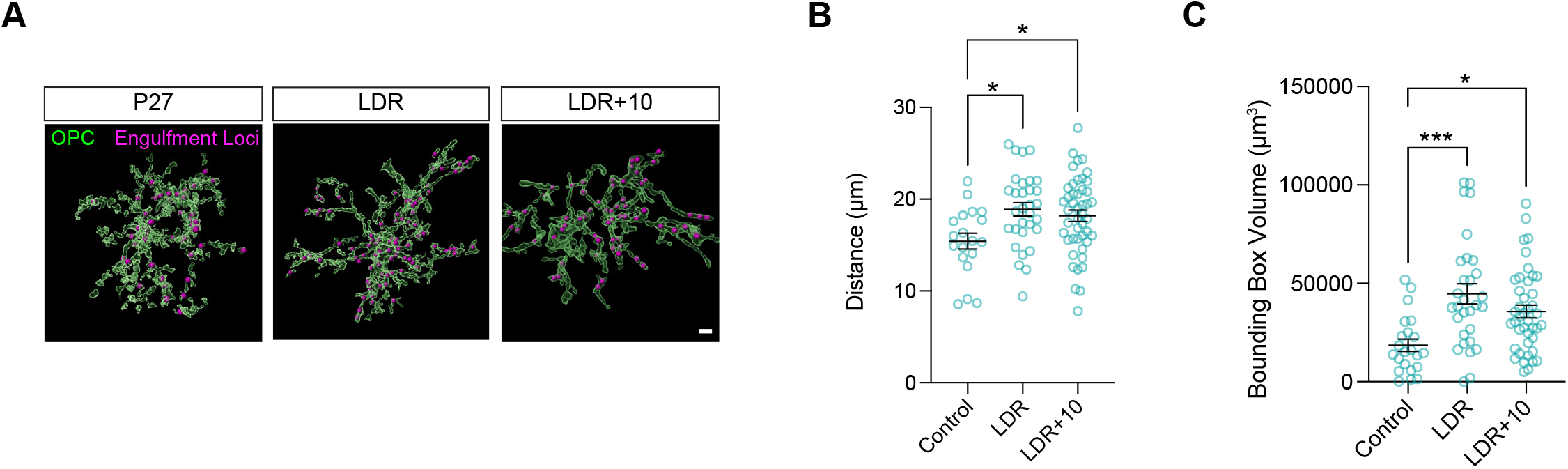
Distribution of engulfment loci across OPCs is regulated by sensory experience. (A) Reconstructions of OPCs (green) and engulfed TC inputs (magenta) illustrating experience-dependent changes in the distribution of points of engulfment across the OPC arbor. Scale bar, 3 μm. (B) Quantification of the distance between the center of the OPC soma and loci at which engulfed inputs reside. One-way ANOVA (p < 0.005) with Tukey’s post-hoc test; n (OPC): P27 = 19, LDR = 31, LDR+10 = 46, from 3 mice per group. (C) Estimated range of synaptic surveillance by a given OPC based upon the bounding box volume calculated from the distribution of engulfment loci as quantified in (B). One-way ANOVA (p < 0.0005) with Tukey’s post-hoc test; n (OPC): P27 = 22, LDR = 30, LDR+10 = 42, from 3 mice per group. (B) and (C) Individual data points with mean +/− SEM. *p< 0.05, ***p<0.001.

